# Gestational diabetes mellitus dysregulates the PD-1/PD-L1 axis at the feto-maternal interface

**DOI:** 10.1101/2023.01.25.525478

**Authors:** ZN. Mihalic, O. Kindler, S. Raftopoulou, A. Santiso, C. Wadsack, A. Heinemann, J. Kargl

**Author notes:** Correspondence to Julia Kargl.

## Abstract

The most common pregnancy complication is gestational diabetes mellitus (GDM), which is a glucose tolerance disorder. Obesity and older maternal age, which are associated with low-grade systemic inflammation, are the main risk factors for GDM. To evaluate the complexity and differences in the immune landscape at the fetal-maternal interface, we examined the maternally derived tissue, decidua basalis (DB), from healthy women, women with obesity, and women with GDM using flow cytometry, western blot, and gene expression analysis. Our results showed that the immune cell composition of DB is not altered by obesity; however, in GDM pregnancies, the DB displays a dysregulated PD-1/PD-L1 axis and significantly reduced regulatory T cell (Treg) infiltration, suggesting reduced local immunosuppression. Our study provides a detailed picture of the immune landscape at the fetal-maternal interface in normal, obese, and GDM pregnancies. This will aid our understanding of possible dysfunctional immune mechanisms in GDM.

## INTRODUCTION

Gestational diabetes mellitus (GDM) is a condition of glucose intolerance that emerges during pregnancy. It is one of the most common pregnancy complications, with an occurrence of up to 20%. Several risk factors contribute to GDM, including age, obesity, ethnicity, and genetics^1^. Obesity has become a major epidemic worldwide^2,3^ and the average maternal age has increased in many countries, leading to increased GDM rates^4,5^. During normal pregnancy, insulin resistance increases, which is compensated by an upregulation of insulin production in pancreatic beta cells. However, in GDM, insulin production is inadequate, which has several short- and long-term consequences for the mother and fetus, such as development of preeclampsia, fetal hypoglycemia after delivery, development of type 2 diabetes for mother and child later in life^6^. The epidemiology, pathophysiology, diagnostics, and prevention of GDM have been extensively reviewed^1,6–9^.

Many studies have shown impaired immune regulation in patients with GDM. Systemic inflammation in GDM is low-grade, with an increase of several inflammatory cytokines in the blood^10,11^. First, studies of the lymphocyte compartment demonstrated an upregulation of maternal checkpoint molecules on blood T cells^12^ and reduced suppressive ability of regulatory T cells (Tregs) from the blood of patients with GDM^13^. A recent study also found enhanced Th17 cell levels in blood from patients with GDM^14^. The expression of natural killer (NK) cell receptors, such as CD16, NKp46, and NKp30, is altered in patients with GDM^15^, and placental villi and extravilli show increased CD16-CD56+ and CD16+ CD56+ NKs^16^. Moreover, myeloid cells show altered immune mechanisms in GDM. Neutrophil blood counts are elevated in the first trimester in women who go on to develop GDM^17^. Additionally, neutrophils isolated from the blood of patients with GDM form more neutrophil extracellular traps (NETs) and affect cytotrophoblasts *in vitro*^18^. GDM is associated with preeclampsia, in which peripheral blood neutrophils are altered and activated. Similar to other blood immune cell populations, blood monocyte counts are elevated in GDM^19^. The role of macrophages in diabetes has been extensively investigated in the placenta. Higher numbers of CD163-expressing macrophages have been found in all compartments of the placenta using immunohistochemistry. Furthermore, higher levels of soluble CD163 have been detected in the circulation of patients with GDM^20,21^. Isolated rat Hofbauer cells (placenta M2 macrophages) switched to a more proinflammatory M1 phenotype when treated with high levels of glucose *in vitro*^22^; however, another study showed that Hofbauer cells retain an immune-regulatory phenotype in GDM placentas^23^.

Systemic inflammation, phenotypes, and function of immune cells in circulation have been well-described in patients with GDM; however, limited data is available on the local immune microenvironment at the fetal-maternal interface, particularly at the decidua basalis (DB). Immune cells are highly abundant in DB and a fine balance between activation, suppression, and regulation of innate and adaptive immune mechanisms is essential for proper placental development, protection, and gas and nutritional exchange within the fetus. Because of the important role of immune cells at the DB, we analyzed the detailed immune landscape in the DB from term pregnancies and compared them with the DB from GDM pregnancies. We hypothesized that the immune response is dysregulated at the DB of GDM pregnancies compared with healthy pregnancies.

In this study, we generated a comprehensive profile of the immune microenvironment using flow cytometry in healthy, obese, and GDM term pregnancies. Our data provide a resource to understand immune cell abundance and interactions, and we identified a dysregulated PD-1/PD-L1 axis and reduced Treg abundance in GDM, leading to decreased immunosuppression.

## RESULTS

### The global immune microenvironment of the DB

GDM is a leading pregnancy complication worldwide, with numerous adverse effects for the mother and the child. The exact mechanism and contribution of immune cells to the progression of GDM and subsequent complications in the placenta remain unknown. In this study, we used high-dimensional flow cytometry together with bioinformatics to study the immune microenvironment at the end of pregnancy at the DB in healthy, obese, and GDM pregnancies (Figure 1a).

**Figure 1.**
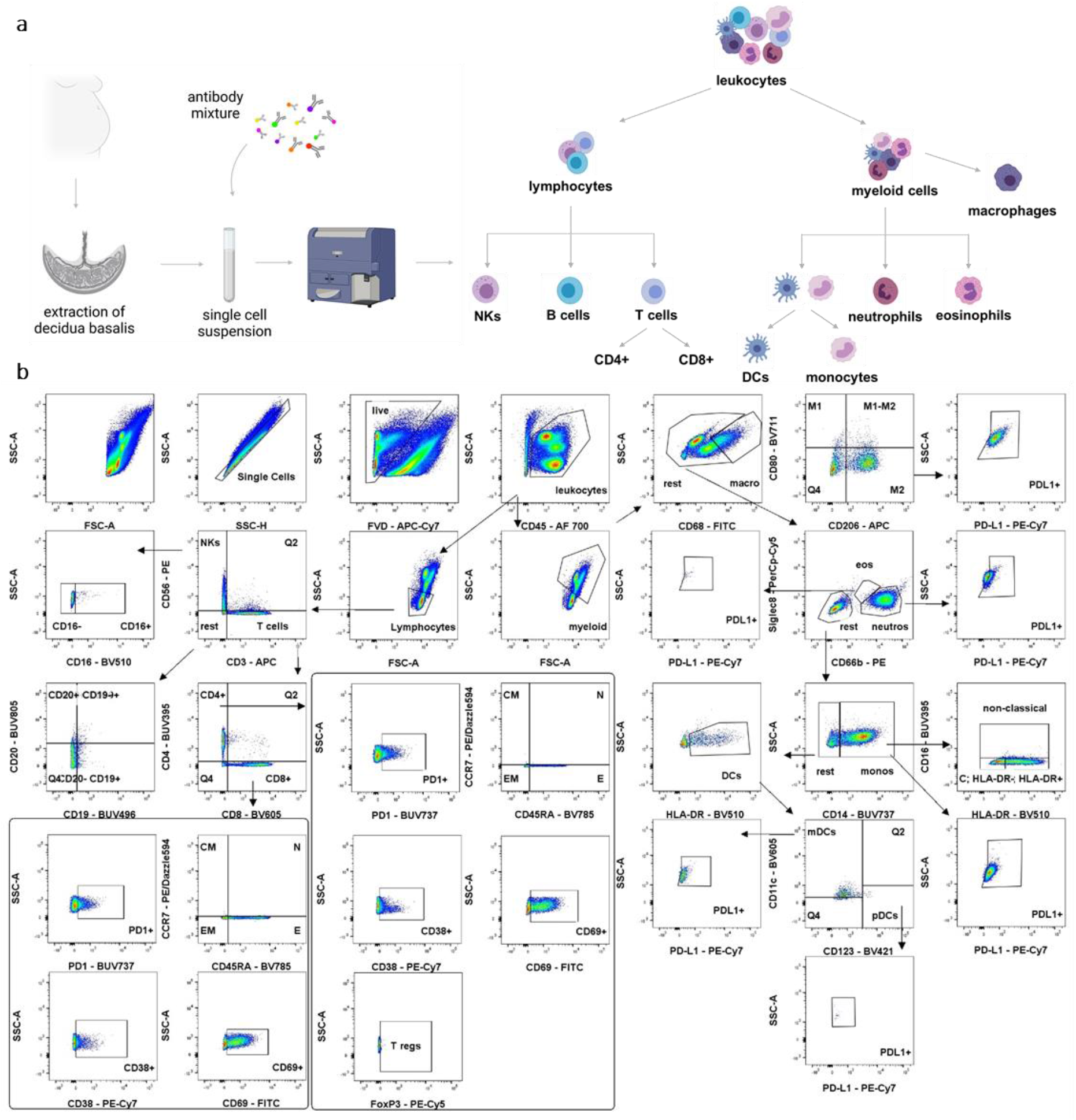
Identification of immune cell types in the DB of term pregnancies. (**a**) Graphical diagram illustrating the experimental workflow. Tissue was processed and a single cell suspension was isolated for flow cytometry analyses, followed by the hierarchical organization of the leukocyte populations. (**b**) Representative dot plots demonstrating the gating strategy to identify immune populations in the DB. The first gate represents all cells, followed by the gate to eliminate doublets. Furthermore, live (FVD) and CD45+ cells were selected. For lymphocyte populations, a size gate was applied, followed by CD3 and CD56 to identify NKs (CD3-, CD56+), B cells (CD3-, CD56-, CD19+) and T cells (CD3+, CD56-). NK subsets were further classified as CD16- or CD16+. T cell subsets were identified as CD4+ and CD8+, and they were further classified as CM, N, E, and EM subsets based on the expression of CD45RA and CCR7. Tregs were defined as CD4+ FoxP3+ T cells. Activation status of CD4+ and CD8+ T cells was determined by expression of PD-1, CD38 and CD69. Myeloid cells were selected by size gating. Subsequently, macrophages were separated as CD68hi, neutrophils as CD66b+, Siglec8-, and monocytes as CD14+. Macrophages were further classified as M1 (CD80+, CD206-), M1-M2 (CD80+, CD206+) and M2 (CD80-, CD206+). Eosinophils were classified as CD66b-, Siglec 8+ cells. Furthermore, DCs were defined as CD14-, HLA-DR+ and subclassified as mDCs (CD11c+, CD123-) and pDCs (CD11c-, CD123+).

A detailed description of patient characteristics is presented in Table 1. GDM was diagnosed by applying an oral glucose tolerance test (oGTT). We included 18 GDM and 24 healthy women in this study regardless of their body mass index (BMI); however, the average maternal age of the GDM group was higher than that of the healthy group. As outlined, maternal age is an independent risk factor in the pathogenesis of GDM (Table 1). BMI of both groups was similar, with an average of 25.1 in normal, healthy women and 25.3 in the GDM group. The normal group was further divided by BMI into three subgroups: BMI <25 (n=13), 25≤ BMI>30 (n=5), and BMI ≥30 (n=6). BMI was also used to subgroup the GDM group: BMI <25 (n=8), 25≤ BMI >30 (n=5) and BMI ≥30 (n=5).

**Table 1.** Clinical parameters of the normal and GDM groups. Student’s t-tests were performed to compare two groups. Data are represented as mean ± SD, *P < 0.05.

|  | *normal* | *GDM white A* | *p value* |
| --- | --- | --- | --- |
| *N* | 24 | 18 |  |
| *BMI* | 25.1 ± 5.5 | 25.3 ± 5.2 | 0.6308 |
| *BMI*<25 | 21.15±2.18 (N=13) | 20.58±1.7 (N=8) | 0,5377 |
| 25≤*BMI*>30 | 27.41±1.54 (N=5) | 26.8±1.53 (N=5) | 0,5484 |
| *BMI*≥30 | 34.08±2.55 (N=6) | 32.63±2.06 (N=5) | 0,3323 |
| *SSW* | 38.9 ± 1.6 | 38.9 ± 2.3 | 0.9642 |
| *placental weight* | 592.9 ± 99 | 605 ± 134.9 | 0.7451 |
| *baby weight* | 3349 ± 468 | 3457.1 ± 346.2 | 0.4257 |
| *age* | 29.6 ± 4.9 | 33.5 ± 5.8 | 0.0267 * |
| *OGGT 0* | 79.33 ± 6.9 | 91.7 ± 9.3 | <0.0001 **** |
| *OGTT 60* | 119.5 ± 21.9 | 168.7 ± 25.6 | <0.0001 **** |
| *OGTT 120* | 99.8 ± 21.5 | 134.1 ± 28.9 | 0.0001 *** |

Flow cytometry panels were applied, consisting of 25 markers (Supplementary Table 1), allowing for the identification of 37 immune populations and subpopulations. Additionally, the activation states of distinct immune populations were determined. The gating strategy for lymphoid and myeloid cells is depicted in Figure 1b. First, we investigated the distribution of major lymphocyte and myeloid cell populations in a representative sample using uniform manifold approximation and projection (UMAP). Based on the median fluorescence intensity expression of the depicted markers, we defined natural killer (NK) (CD56+) and T cell (CD3+) clusters among the lymphocyte populations. T cells were separated into helper (CD4+) and cytotoxic (CD8+) T cells. The main myeloid clusters include neutrophils (CD66b+Siglec8-), monocytes (CD14+), and macrophages (CD68+). Macrophages were further separated into M1 (CD206-CD80+) and M2 (CD206+CD80-) subsets (Figure 2a). When comparing lymphocyte populations of the samples, we observed the highest infiltration of T cells within the DB, of which CD4+ T cells represented the highest proportion. By examining the expression of CCR7 and CD45RA on the T cell surface, we classified CD4+ and CD8+ T cells into effector memory (EM), effector (E), central memory (CM), and naïve (N) subpopulations. EM cells made up a major subgroup of CD4+ and CD8+ T cells, as well as E cells, while CM and N CD4+ or CD8+ T cells represented minor subpopulations. As previously reported, regulatory NK cells (CD56+CD16-) were the dominant NK subpopulation^24^. CD19+CD20-B cells represented the smallest lymphocyte population.

**Figure 2.**
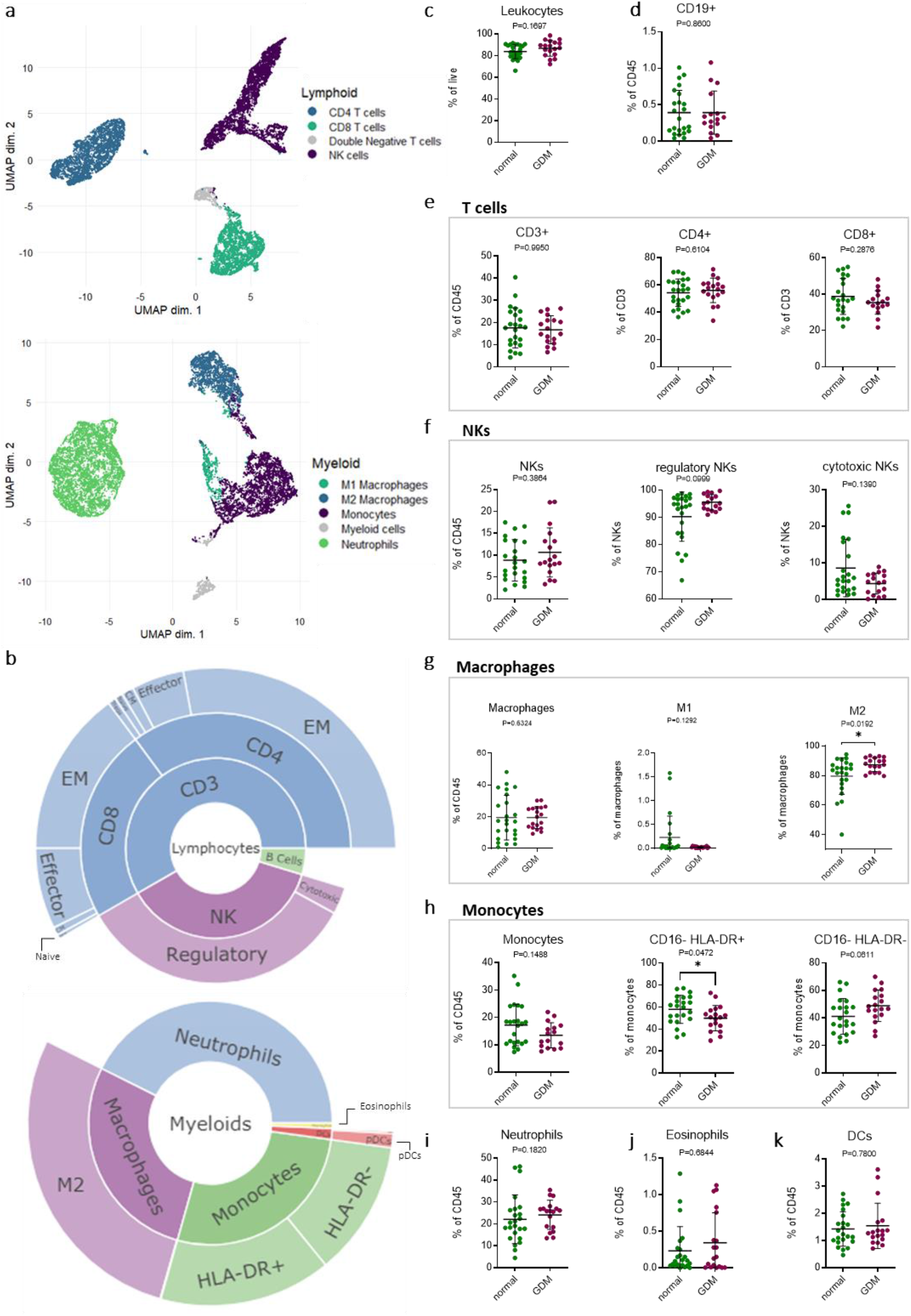
Changes in immune populations in the DB of normal and GDM tissue. (**a**) Representative UMAP plots of the lymphocyte and myeloid populations. (**b**) Sunburst plots showing the average distribution of immune populations of the lymphocyte and myeloid cell lineages of all samples; effector memory (EM), effector (E), central memory (CM), naïve (N). (**c–k**) Abundance and changes in the main immune populations between normal (N = 24, left) and GDM (N = 18, right) DB samples. CD19+ B cells, T cells, NKs, macrophages, monocytes, neutrophils, eosinophils, and DCs are represented as percentages of CD45+ cells. Subpopulations of T cells, NKs, macrophages, and monocytes are represented as a percentage of the parent population. Mann–Whitney U tests were performed to compare two groups. Data are represented as mean ±SD, *P < 0.05.

In the myeloid compartment, neutrophils represented more than one-third of all immune cells, followed by monocytes and macrophages in approximately equal amounts. The majority of macrophages in the DB were M2 macrophages, which has previously been reported^25^. Furthermore, half of the monocytes expressed HLA-DR, a surface protein important for antigen presentation.

Eosinophils and dendritic cells (DCs) were the smallest populations in the myeloid lineage, with a majority of DCs being myeloid-derived DCs (Figure 2b).

### The main immune populations of the DB do not show major differences between controls and subjects with GDM

We observed high leukocyte infiltration in the DB, with CD45+ cells representing 66%–99% of total live cells, both in healthy and GDM pregnancies (Figure 2c). This is in part due to our *ex vivo* cell isolation technique, which enriches for CD45+ leukocytes, allowing for further immunologic analysis in this study. Infiltration of B cells was very low in the DB, making up a total of 1% of all leukocytes in both groups (Figure 2d, Supplementary Figure 1b). T cells represented 20% of all leukocytes, of which 60% were CD4+ T cells and 40% were CD8+ T cells. There was no difference in the infiltration of these cell types in GDM pregnancies compared to healthy pregnancies (Figure 2e). CD4+ and CD8+ cells were further defined as CM, E, EM, and N cells, but no differences between the healthy and GDM groups were observed (Supplementary Figure 1a). Approximately 10% of leukocytes represent NKs as CD56+CD3-cells, which were further divided into regulatory and cytotoxic NK cells based on the expression of CD16. More than 90% of all NK cells were regulatory cells in the DB, which correlates with previously published data (Figure 2f)^24^. In the myeloid lineage, macrophages are one of the most prevalent populations together with neutrophils and monocytes, each representing about 20% (Figure g-i). M2 macrophage polarization was significantly increased in GDM pregnancies; however, in both groups, >90% of all macrophages were of the M2 phenotype. Approximately half of the monocytes expressed HLA-DR, with a lower level of HLA-DR measured in GDM individuals compared with normal (Figure 2h). Additionally, we did not observe differences in eosinophils and DC infiltration between the two groups (Figure 2j–k, Supplementary Figure 1c).

### The PD-L1/PD-1/Treg pathway is dysregulated in GDM pregnancies

The expression of immune checkpoint molecules is essential for immune cell activation and function. We therefore evaluated the activation state of lymphocyte populations via the expression of CD69, CD38, and PD-1 and the expression of PD-L1 in myeloid populations. Furthermore, Tregs play a major role in normal pregnancy in the first trimester, and their dysregulation contributes to the development of pregnancy complications^13^. Principal component analysis (PCA) of the total composition of lymphocyte populations did not reveal major differences between healthy and GDM pregnancies (Figure 3a, left panel). However, myeloid populations were distinct, with expression of the PD-L1 checkpoint molecule as a major factor driving separation between the healthy and GDM groups (Figure 3a, right panel). Therefore, we evaluated the expression of PD-L1 on live cells and observed a downregulation of total PD-L1 in the GDM group. Further analysis of PD-L1 expression on immune (CD45+) and non-immune (CD45-) cells revealed that non-immune cells (e.g. extravillous trophoblasts EVTs) show high expression of PD-L1 but with no difference between healthy and GDM samples. High expression of PD-L1 on EVTs has been previously reported^26^. Contrastly, there was a significant decrease in expression of PD-L1 on CD45+ DB leukocytes from individuals with GDM (Figure 3b). Furthermore, we analyzed the expression of the activation molecules PD-1, CD38, and CD69 in lymphocyte populations and PD-L1 in myeloid populations. PD-L1 expression was reduced in macrophages and monocytes from GDM patients, but no differences were found in neutrophils (Figure 3c, Supplementary Figure 2). Consistent with these results, we observed downregulation of PD-1 on CD8+ T cells and a trend towards decreased expression in CD4+ T cells (Figure 3d). A heatmap displaying expression of PD-1 and PD-L1 for each individual sample demonstrated an overall reduction of these two surface markers in GDM patients compared with healthy (Figure 3e). In line with previous studies on peripheral blood Tregs, we observed lower Treg infiltration in DB from GDM samples (Figure 3f)^14^. The T cell activation markers CD38 and CD69 did not differ between healthy and GDM DB (Figure 3g). To test whether the expression of PD-L1 was lower at transcriptional and translational levels, we performed quantitative PCR (qPCR) and western blotting (WB) of tissue homogenates to measure PD-L1 mRNA and protein levels, respectively. qPCR and WB did not show significant differences between the samples, possibly due to small sample size (n=5-7/group). However, we did observe a trend toward lower mRNA and protein PD-L1 expression in tissue homogenates (Supplementary Figure 3). Since GDM is determined using an oGTT, we investigated whether Treg infiltration and PD-L1 expression on macrophages and monocytes correlate with higher maternal glucose levels after a 60-min oGTT. Treg infiltration and PD-L1 levels of monocytes negatively correlated with higher glucose levels, and PD-L1 levels in macrophages showed a trending negative correlation with higher glucose levels (Figure 3h). Upon analysis of PD-L1 expression and infiltration of Tregs using publically available, previously published scRNAseq data, we confirmed expression of PD-L1 on macrophages and EVTs (Supplementary Figure 4)^27^.

**Figure 3.**
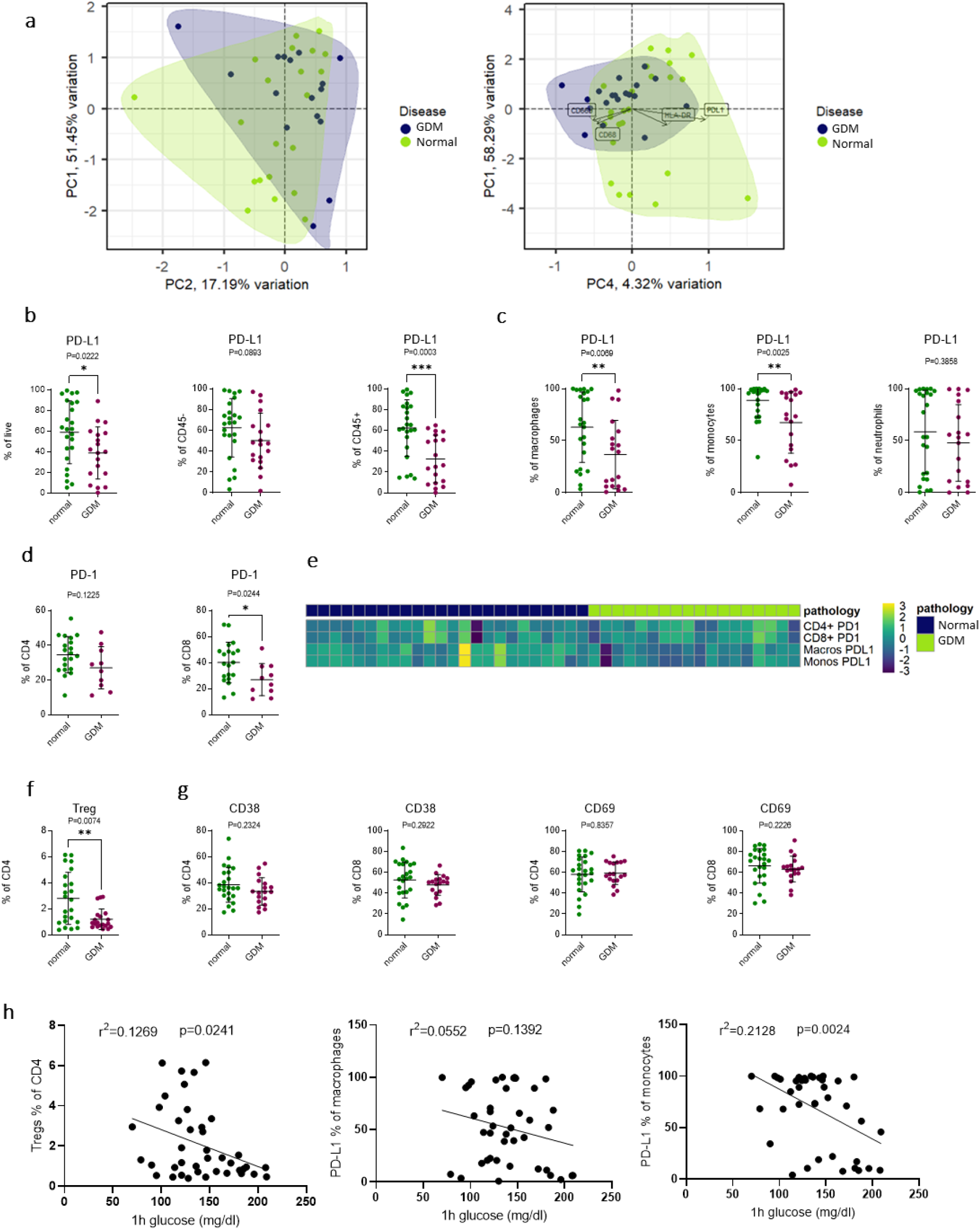
PD-L1/PD-1/Treg axis is downregulated in GDM pregnancies. (**a**) Principal component analysis of flow cytometry data of the lymphocyte (left) and myeloid (right) populations separating normal and GDM samples. (**b**) Abundance and changes in the PD-L1 expression of live, CD45-, and CD45+ cells. (**c)** Abundance and changes in PD-L1 expression between normal (N = 24, left) and GDM (N = 18, right) samples in neutrophils, monocytes, and macrophages. (**d**) Abundance and changes of PD-1 expression between normal (N = 24, left) and GDM (N = 18, right) samples on CD4+ and CD8+ T cells. (**e**) Heatmap representation presenting sample-to-sample variability in the expression of PD-L1 and PD-1 checkpoint molecules. (**f**) Treg cells displayed as a percentage of CD4+ T cells. (**g**) Abundance and changes in stimulatory molecule expression between normal (N = 24, left) and GDM (N = 18, right) samples on CD4+ and CD8+ T cells. (**h**) Linear correlation of Treg infiltration (P = 0.0241, r2 = 0.1269, N =41) and PD-L1 expression (P = 0.01392, r2 = 0.0552, N = 41) with oGTT after 60 min. Mann–Whitney U tests were performed to compare two groups. Data are represented as mean ± SD, *P < 0.05.

### BMI does not influence immune changes in normal donors

Maternal obesity is one of the biggest risk factors for GDM development. Therefore, we next investigated how maternal obesity influences immune cell composition at the DB and whether the changes we previously observed in patients with GDM are driven by higher maternal BMI. We divided normal pregnancies into three groups using a standard classification model^28^ based on the BMI of the mother before pregnancy: BMI < 24.9 was considered normal, a BMI of 25–29.9 was considered overweight, and a BMI > 30 was considered obese. We did not observe differences among the main immune populations in these BMI groups (Figure 4a). Furthermore, when investigating immune subpopulations, no differences among CD4+ or CD8+ T cell subpopulations, CD20+ B cells, and DCs were observed (Figure 4a and supplementary Figure 5). Again, we evaluated expression of functional markers on immune cells and observed no differences (Figure 4b-e and supplementary Figure 5). Altogether, these results demonstrate that immune cell composition of the DB during pregnancy is not correlated with or driven by BMI.

**Figure 4.**
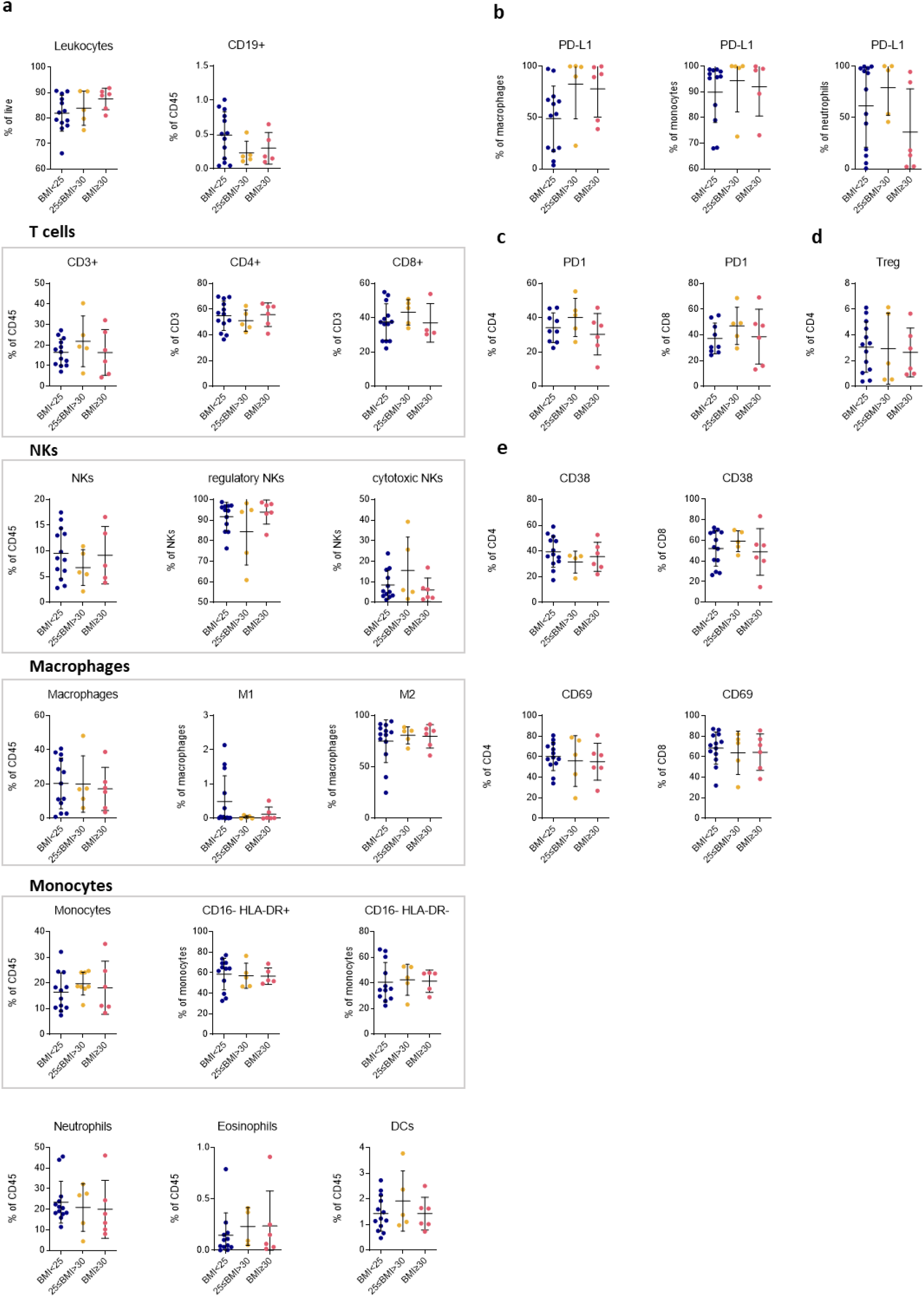
BMI does not influence immune changes in normal donors. (**a**) Abundance and changes in the main immune populations between normal DB samples from normal BMI (N=13), overweight (N=5), and obese mothers (N=6). CD19+, T cells, NKs, macrophages, monocytes, neutrophils, eosinophils, and DCs are represented as the percentage of CD45+ cells. Subpopulations of T cells, NKs, macrophages, and monocytes are represented as percentages of the parent populations. (**b**) Abundance and changes in PD-1 expression between normal DB samples from normal BMI, overweight, and obese mothers. (**c**) Abundance and changes in PD-L1 expression between normal DB samples from normal BMI, overweight, and obese mothers on neutrophils, monocytes, and macrophages. (**d**) Treg cells displayed as a percentage of CD4+ T cells. (**e**) Abundance and changes of stimulatory molecule expression in normal DB samples from normal BMI, overweight, and obese samples on CD4+ and CD8+ T cells. One-way ANOVA was performed to compare three groups. Data are represented as mean ± SD, *P < 0.05.

## DISCUSSION

Decidual immune cells are one of the main players in the development and maintenance of maternal immune tolerance toward the fetus. Here, we present a detailed examination of the immune cell landscape in the decidua basalis at term. Most studies of immune involvement in metabolic deranged pregnancies, as in GDM, have been conducted on peripheral blood immune cells and not at the local fetal-maternal tissue interface. Therefore, we have comprehensively characterized the immune microenvironment in the DB of women with GDM compared with normal, non-GDM pregnancies. Here, we showed that leukocytes infiltrate in into the DB of both normal and GDM individuals. Other cell types have also been reported to infiltrate the DB, such as populations of trophoblasts, stromal cells, endothelial cells, and others^29^. We did not observe differences in the main immune populations comparing normal and GDM pregnancies. The B cell population is one of the smallest immune cell populations in the DB, followed by eosinophils and DCs. Previously, an increase in B cells in the peripheral blood was found to correlate with insulin resistance^30^, although we did not observe similar differences in the DB. NK cells are a major immune population, defined as CD56+ CD16-cells in the decidua of the uterus, and they produce chemokines and growth factors important for normal EVT function and invasion^24^. Moreover, according to a previous study, decidua NKs cannot polarize microtubule-organizing centers and perforin-containing granules to the synapse due to their lack of cytotoxic properties^31^. This correlates with our data, confirming that a very small subset expresses CD16+. Previously, CD16+ expression was reported to be increased on peripheral blood NK cells, and low-grade inflammation was observed in GDM^15^. Transcriptomic profiling with scRNAseq of two GDM placental samples revealed increased NK infiltration in the placenta and upregulation of CD16 expression^27^. In our study, we did not observe differences in NK subsets comparing GDM and normal DB. Therefore, additional research should focus on the phenotype of NKs in GDM at the DB. At the end of pregnancy, a reduction in NK cells and upregulated infiltration of T cells in the DB have been observed^32^.

Many studies have investigated the role of T cells in pregnancy, with a focus on Tregs. It has been shown that maternal T cells can recognize fetal alloantigens, and in mismatched pregnancies CD4+ cells are activated^33^. Correlating with our data, it has been previously found that the biggest subset of CD4 and CD8 T cells are EM^34^. In the blood of GDM subjects, an increase of CD4+ cells with a proinflammatory phenotype has been described^14^. Our data did not show significant differences in the infiltration of CD4 and CD8 T cells locally in the DB.

Macrophages contribute to normal EVT invasion and tissue remodeling during pregnancy^25^. As previously reported, most macrophages in the decidua have an M2 phenotype at the end of pregnancy. In the placenta, Hofbauer cells have increased expression of CD206 in patients with GDM^23^. However, one study showed that Hofbauer cells switched toward an M1 phenotype after treatment with high glucose levels *in vitro*^22^. Additionally, higher macrophage levels have been reported in the placentas of patients with GDM^35^. At the DB, we did not observe significant differences in the number of macrophages; however, there was a slight shift toward an anti-inflammatory phenotype, which could represent a compensatory mechanism for maintaining normal immune tolerance at the DB. Blood monocytes have mostly been researched in the context of pregnancy versus locally at the DB, which could account for these contrasting results. There are lower numbers of classical monocytes in the blood of pregnant women when compared with nonpregnant women, with a decrease in preeclampsia and GDM^19,36^. Expression of HLA-DR on the surface of monocytes is responsible for antigen presentation to T cells, and HLA-DR expression is associated with an inflammatory state^37^. Here, we report high expression of HLA-DR on monocytes in the DB. We observed a slight decrease in expression in patients with GDM and a trending increase in monocytes that do not express HLA-DR. However, infiltration of neutrophils, eosinophils, and DCs was similar in normal versus GDM samples.

Furthermore, we observed downregulation of the PD-L1/PD-1/Treg pathway in DB from GDM individuals versus healthy. Tregs are a subpopulation of CD4+ T cells with immunosuppressive properties. Treg suppression of maternal immune mechanisms at the DB is a necessary mechanism for fetal development^38^. Recently, DB Tregs were shown to specifically recognize matched human leukocyte antigen-C, showing T cell activation in mismatched pregnancies^33^. Additionally, uterine Tregs show a higher activation state with a similar transcriptional signal as Tregs from tumors^39^. The PD-L1/PD-1 axis has been a focus of cancer research, and PD-1 and PD-L1 blocking antibodies are used as successful antitumor therapies. The PD-L1 molecule, expressed on different cells including myeloid and nonimmune cells, is the ligand for PD-1 and is expressed by lymphocytes^40^. Higher PD-L1 expression on Tregs correlates with higher suppression properties^41^. Similarly, blockade of PD-L1 resulted in reduced Treg expansion in mouse studies^42^. Even more important, the PD-L1/PD-1 pathway is not only important in cancer treatment but also in the development of type 1 diabetes. Antitumor treatment with immune checkpoint inhibitors, including PD-L1 and PD-1 blockade, is associated with the induction of autoimmune diabetes^43^. Thus, the PD-1/PD-L1 pathway may influence T cell exhaustion in the pancreas, leading to T cell-mediated destruction of pancreatic ß cells^44^. Decreased PD-1 and PD-L1 expression was measured in recurrent miscarriage samples^45^. In the serum of pregnant women, increased soluble PD-L1 levels were found, suggesting the suppression of the maternal immune response^46^. PD-L1 treatment *in vitro* drove M1 macrophage differentiation, and PD-1 treatment increased phagocytic activity^45^. Furthermore, depletion of Tregs abolished the PD-L1 blockade effect, and Treg transfer into PD-L1-depleted mice improved fetal survival^47^. In preeclampsia, the PD-1/PD-L1 pathway is altered, leading to the dysregulation of Treg responses^48^. Thus, the PD-L1/PD-1/Treg pathway is extremely important for normal immune regulation at the DB. In GDM pregnancies, Tregs isolated from blood have a reduced suppressive activity compared with healthy controls^13^. A study showed that Rank depletion in the thymic epithelium causes inadequate Treg accumulation in the placenta, which also resulted in GDM development^49^. PD-1 has been shown to be downregulated on blood T cells in patients with GDM^50^. Similarly, we observed this pattern at the DB. Dysregulation of the PD-1/PD-L1 pathway may be an important factor in the development of pregnancy complications, including GDM, and the immune cell imbalances reported here may lead to lower immune suppression in the DB. Conversely, obesity does not seem to affect the expression of checkpoint inhibitors or the infiltration of Tregs. Thus far, several cytokines have been associated with GDM. Both upregulation (CCL2, CXCL1, CXCL8, CXCL9, and CXCL12) and downregulation (CCL4, CCL11, and CXCL10) of cytokines and chemokines has been reported in the blood of patients with GDM. These signaling molecules affect immune responses in the pancreas. Therefore, more research must be conducted to investigate the role of these cytokines in GDM and at the DB^11^.

In conclusion, the prevalence of GDM is rising and becoming one of the most common health complications during pregnancy. The immune microenvironment plays an important role in GDM pathology. However, most research has focused on peripheral immune cells. Thus, we hereby present a comprehensive study of the DB immune composition in normal and GDM pregnancies. We observed a disbalanced PD-1/PD-L1/Treg pathway in GDM samples. Therefore, more research should focus on immune mechanisms at the DB in pregnancy complications and investigate the exact immunoregulatory mechanisms of the PD-1/PD-L1/Treg pathway in GDM.

## METHODS

### Study design

Study participants were recruited from the Department of Obstetrics and Gynaecology, Medical University of Graz (Graz, Austria) (Table 1). Informed consent was obtained from all participants. Placenta tissue samples were collected from singleton pregnancies and processed within 4 h after delivery by cesarean section or vaginal delivery. GDM was diagnosed by oGTT within the second trimester of pregnancy according to WHO criteria as outlined here: Fasting plasma glucose ≥ 92 mg/dl, a 1-h plasma glucose ≥ 180 mg/dl, or a 2-h plasma glucose ≥ 153 mg/dl^51^.

### Study approval

This study complied with the Declaration of Helsinki and was approved by the Ethics Committee of the Medical University of Graz (EK-number: 29-319 ex 16/17).

### Tissue processing

Tissues were received within 4 h after delivery and immediately processed and stained for flow cytometry analyses as previously described^52^. In brief, the DB was carefully separated from the placenta, as previously described,^53^ and washed twice in 40 ml PBS without Ca^2+^ and Mg^2+^ for 2 min while shaking. Subsequently, the remaining villous part was removed from the tissue, and the tissue was washed again to remove any remaining blood. Small pieces were snap-frozen for further experiments. The decidua was mechanically dissociated and digested in RPMI-1640 supplemented with 40 U ml^-1^ DNAse I (Worthington) and 300 U ml^-1^ collagenase I (Worthington) at 37°C for 30 min. The tissue digest was homogenized by passing through a 19-G needle and a 100-μm strainer. Digestion was stopped with the addition of fetal bovine serum (FBS). Erythrocytes were lysed using 1× RBC lysis buffer (BioLegend). Lysis was inactivated by adding four volumes of PBS without Ca^2+^ and Mg^2+^. Finally, the decidual cells were passed through a 50-μm strainer. Cell counts and viability were determined using an EVE automated cell counter (NanoEntek).

### Flow cytometry

Viability staining was performed immediately after forming a single cell suspension, and staining was performed in three separate panels. The cells were stained with fixable viability dye (FVD) eFluorTM 780 (eBioscience) before fixation.

#### Lymphocyte surface panel

The cells were fixed for 10 min in IC fixation buffer (Invitrogen), washed in staining buffer (SB, PBS without Ca^2+^ and Mg^2+^ supplemented with 2% FBS), and stained the following day. First, the cells were preincubated with Fc receptor blocking solution (Biolegend) to reduce nonspecific binding and subsequently stained with fluorochrome-conjugated anti-human antibodies against CD45, CD3, CD4, CD8, CCR7, CD45RA, CD38, CD69, PD-1, CD19, CD20, CD56, and CD16 (for antibody and clone details, see Supplementary Table 1) for 30 min at 4°C and washed twice in SB.

#### Lymphocyte intranuclear panel

The cells were fixed for 10 min in 1× TF fixation buffer (BD), washed in SB, and stained the following day. Before staining, the cells were again fixed and permeabilized for 40 min using a BD transcription factor fixation/permeabilization kit to allow antibodies to bind to nuclear proteins. Subsequently, the cells were preincubated with Fc receptor blocking solution and fluorochrome-conjugated anti-human antibodies against CD45, CD3, CD4, and FoxP3 (for antibody and clone details, see Supplementary Table 1) for 30 min at 4°C and washed twice in TF washing buffer (BD) and once in SB.

#### Myeloid intracellular panel

The cells were fixed for 10 min with the BD Cytofix kit, washed in BD Cytofix wash, and stained the following day. Prior to staining, the cells were again fixed and permeabilized for 20 min using the BD Cytofix/Cytoperm kit. Subsequently, the cells were preincubated with Fc receptor blocking solution and fluorochrome-conjugated anti-human antibodies against CD45, CD68, CD80, CD206, CD66b, Siglec 8, CD14, HLA-DR, CD16, CD11c, CD123, and PD-L1 (for antibody and clone details, see Supplementary Table 1) for 30 min at 4°C and washed twice in Cytofix wash (BD) and once in SB.

#### Analysis

All samples were analyzed on a BD LRSFortessa flow cytometer with FACSDiva software (BD). Compensation was performed using single stains, and cut-offs for background fluorescence were based on the fluorescence minus one (FMO) strategy. Gating for each sample was based on an SSC-Height versus SSC-Area plot to eliminate aggregates. FVD staining was used to identify and eliminate dead cells. Analysis and compensation were performed using FlowJo software (TreeStar). For PCA plots, fluorescence intensities were exported to an xlsx-file, imported into R (v.4.2.1) with the readxl package, and arcsinh-transformed with a cofactor of 150. PCA plots were drawn with PCAtools (https://github.com/kevinblighe/PCAtools). Heatmaps were plotted with the pheatmap package (https://cran.r-project.org/web/packages/pheatmap/index.html), and colors were used from the viridis package (https://cran.r-project.org/web/packages/viridis/viridis.pdf). UMAP plots were drawn with the cytofWorkflow package for a representative patient (https://github.com/markrobinsonuzh/cytofWorkflow). For the sunburst plots, the percentages of cell types were rounded, and the plotly package was used^54^.

### scRNAseq analysis

A previously published dataset was used for analysis^27^. Unique molecular identifier count matrixes were download via the GEOquery package (2.58.0) from GSE173193 and processed with the R package Seurat (v.4.0.6). Low-quality cells (unique genes detected at <400 and >4000 per cell and >10% mitochondrial genes) were excluded. For batch correction, the R package harmony (v.0.1.0) was used (https://cran.r-project.org/web/packages/harmony/index.html). The resulting dimensions were used for clustering, and the cluster identities were assigned using expression markers from the original analysis. Cluster annotation was additionally verified with sctype (R version 4.0.5 (2021-03-31); Platform: x86_64-conda-linux-gnu (64-bit); Running under: Debian GNU/Linux 11 (bullseye))^54–56^.

### Quantitative PCR

Total RNA was extracted from frozen tissue samples using ceramic beads (Precellys homogenizer, 1× 20 s) in RLT buffer (RNeasy Mini kit, Qiagen) supplemented with 1%ß-mercaptoethanol. Subsequently, RNA was extracted following the manufacturer’s protocol (Qiagen), and the RNA concentration was measured with Nanodrop. The RNA concentration of the samples was adjusted to the RNA concentration of the sample with the lowest RNA content with RNAse-free water. Reverse transcription was performed using 10 μl reverse transcription master mix (Applied Biosystems) mixed with 10 μl RNA in a Thermal Cycler (BioRad) for 10 min at 25°C, 120 min at 37°C, 5 min at 85°C, and cooled down to 4°C. 9 μl TaqMan Gene Expression Master Mix (Applied Biosystems) was used along with 1 μl PD-L1 (Hs00204257_m1, FAM, Applied Biosystems) and 1 μl GAPDH (Hs4325792_m1, VIC, Applied Biosystems) probes mixed with 9 μl cDNA. Real time PCR was performed using the CFX Connect Real Time System (BioRad) for 2 min at 50°C, 10 min at 95°C, 15 seconds at 95°C, and 1 min at 60°C for 40 cycles. Expression of PD-L1 is presented as the mean of the dCT values of normal and GDM pregnancies.

### Western blot

Total protein was isolated from frozen tissue samples using ceramic beads (Precellys homogenizer, 1x 20 s) in IP buffer (0.1% Triton-X; 150 mM NaCl; 25 mM KCl; 10 mM Tris HCl, pH 7.4; 1 mM CaCl2 in H2O) supplemented with 1:100 protease/phosphatase inhibitor (Cell Signaling). The protein content was measured using a Pierce BCA assay kit (Thermo Scientific). Samples were mixed with 4× NuPage sample buffer (90 μl 4× LDS sample buffer + 10 μl reducing reagent) and boiled for 10 min at 95°C. 20 μg of protein from each homogenate was applied to the NuPage, Bis–Tris gel (Invitrogen) and run for 45 min at 200 V. Subsequently, proteins were transferred to the membrane using the iBlot transfer device. The membrane was blocked in TBST (1× TBS + 0.1% Tween) supplemented with 5% milk for 1 h. The membrane was incubated in the primary antibody mix overnight at 4°C. Anti-PD-L1 (clone E1L3N, 1:1000 dilution, Cell Signaling) and anti-GAPDH (clone 14C10, 1:500 dilution, Cell Signaling) monoclonal antibodies were used. After primary antibody incubation, the membranes were washed for 30 min and subsequently incubated in secondary antibody HRP-Goat anti-rabbit IgG (polyclonal, 1:5000 dilution, Jackson Immunoresearch) for 2h at RT. The membranes were again washed and developed using ECL solutions (BioRad).

### Statistical analysis

GraphPad Prism 9.1 (GraphPad Software) and R software were used to perform statistical analyses. Statistically significant differences between two groups were determined using the Mann–Whitney U test. To compare more than two groups, one-way ANOVA followed by the Turkey’s post hoc test was used. To study the relationship between two variables, linear regression was used. A P-value <0.05 was considered statistically significant.

## Author contributions

ZN.M designed and performed the experiments, analyzed and interpreted the results, designed the figures, and wrote manuscript. O.K. performed the bioinformatics evaluation of the data. S.R. and A.S. assisted with the performance of the experiments. C.W. and A.H. supplied ethical permission and samples and reviewed the manuscript. J.K. designed and supervised the study, interpreted the data, and wrote the manuscript. All authors critically reviewed the manuscript and approved the submitted version.

## Acknowledgments

We are grateful to Sabine Kern, Iris Red, and the study nurses at the Department of Obstetrics and Gynaecology, Medical University of Graz (Graz, Austria), for their excellent technical assistance and Marah Runtsch for proof reading the manuscript.

## REFERENCES

1. McIntyre, H. D. et al. Gestational diabetes mellitus. Nat. Rev. Dis. Primer 5, 47 (2019).

2. Wong, J. C., O’Neill, S., Beck, B. R., Forwood, M. R. & Khoo, S. K. Comparison of obesity and metabolic syndrome prevalence using fat mass index, body mass index and percentage body fat. PloS One 16, e0245436 (2021).

3. Williams, E. P., Mesidor, M., Winters, K., Dubbert, P. M. & Wyatt, S. B. Overweight and Obesity: Prevalence, Consequences, and Causes of a Growing Public Health Problem. Curr. Obes. Rep. 4, 363–370 (2015).

4. Vandekerckhove, M., Guignard, M., Civadier, M.-S., Benachi, A. & Bouyer, J. Impact of maternal age on obstetric and neonatal morbidity: a retrospective cohort study. BMC Pregnancy Childbirth 21, 732 (2021).

5. Breart, G. Delayed childbearing. Eur. J. Obstet. Gynecol. Reprod. Biol. 75, 71–73 (1997).

6. Baz, B., Riveline, J.-P. & Gautier, J.-F. ENDOCRINOLOGY OF PREGNANCY: Gestational diabetes mellitus: definition, aetiological and clinical aspects. Eur. J. Endocrinol. 174, R43–R51 (2016).

7. Plows, J., Stanley, J., Baker, P., Reynolds, C. & Vickers, M. The Pathophysiology of Gestational Diabetes Mellitus. Int. J. Mol. Sci. 19, 3342 (2018).

8. Sweeting, A., Wong, J., Murphy, H. R. & Ross, G. P. A Clinical Update on Gestational Diabetes Mellitus. Endocr. Rev. bnac003 (2022) doi:10.1210/endrev/bnac003.

9. Nguyen-Ngo, C., Jayabalan, N., Salomon, C. & Lappas, M. Molecular pathways disrupted by gestational diabetes mellitus. J. Mol. Endocrinol. 63, R51–R72 (2019).

10. Atègbo, J.-M. et al. Modulation of Adipokines and Cytokines in Gestational Diabetes and Macrosomia. J. Clin. Endocrinol. Metab. 91, 4137–4143 (2006).

11. Liu, H. et al. Chemokines in Gestational Diabetes Mellitus. Front. Immunol. 13, 705852 (2022).

12. Pendeloski, K. P. T. et al. Immunoregulatory molecules in patients with gestational diabetes mellitus. Endocrine 50, 99–109 (2015).

13. Schober, L. et al. The role of regulatory T cell (Treg) subsets in gestational diabetes mellitus. Clin. Exp. Immunol. 177, 76–85 (2014).

14. Sheu, A. et al. A proinflammatory CD4+ T cell phenotype in gestational diabetes mellitus. Diabetologia 61, 1633–1643 (2018).

15. Chiba, H. et al. Expression of Natural Cytotoxicity Receptors on and Intracellular Cytokine Production by NK Cells in Women with Gestational Diabetes Mellitus. Am. J. Reprod. Immunol. 75, 529–538 (2016).

16. Hara, C. de C. P. et al. Characterization of Natural Killer Cells and Cytokines in Maternal Placenta and Fetus of Diabetic Mothers. J. Immunol. Res. 2016, 7154524 (2016).

17. Sun, T. et al. Elevated First-Trimester Neutrophil Count Is Closely Associated With the Development of Maternal Gestational Diabetes Mellitus and Adverse Pregnancy Outcomes. Diabetes 69, 1401–1410 (2020).

18. Stoikou, M. et al. Gestational Diabetes Mellitus Is Associated with Altered Neutrophil Activity. Front. Immunol. 8, 702 (2017).

19. Angelo, A. G. S. et al. Monocyte profile in peripheral blood of gestational diabetes mellitus patients. Cytokine 107, 79–84 (2018).

20. Barke, T. L. et al. Gestational diabetes mellitus is associated with increased CD163 expression and iron storage in the placenta. Am. J. Reprod. Immunol. 80, e13020 (2018).

21. Bari, M. F. et al. Elevated Soluble CD163 in Gestational Diabetes Mellitus: Secretion from Human Placenta and Adipose Tissue. PLoS ONE 9, e101327 (2014).

22. Sisino, G. et al. Diabetes during pregnancy influences Hofbauer cells, a subtype of placental macrophages, to acquire a pro-inflammatory phenotype. Biochim. Biophys. Acta 1832, 1959–1968 (2013).

23. Schliefsteiner, C. et al. Human Placental Hofbauer Cells Maintain an Anti-inflammatory M2 Phenotype despite the Presence of Gestational Diabetes Mellitus. Front. Immunol. 8, 888 (2017).

24. Hanna, J. et al. Decidual NK cells regulate key developmental processes at the human fetal-maternal interface. Nat. Med. 12, 1065–1074 (2006).

25. Svensson, J. et al. Macrophages at the fetal-maternal interface express markers of alternative activation and are induced by M-CSF and IL-10. J. Immunol. Baltim. Md 1950 187, 3671–3682 (2011).

26. Zhang, R. et al. PD-L1 enhances migration and invasion of trophoblasts by upregulating ARHGDIB via transcription factor PU.1. Cell Death Discov. 8, 395 (2022).

27. Yang, Y. et al. Transcriptomic Profiling of Human Placenta in Gestational Diabetes Mellitus at the Single-Cell Level. Front. Endocrinol. 12, 679582 (2021).

28. NIH. Classification of Overweight and Obesity by BMI, Waist Circumference, and Associated Disease Risks (https://www.nhlbi.nih.gov/health/educational/lose_wt/BMI/bmi_dis) Accessed on 05.09.2022 11:40.

29. Vento-Tormo, R. et al. Single-cell reconstruction of the early maternal–fetal interface in humans. Nature 563, 347–353 (2018).

30. Zhuang, Y. et al. B Lymphocytes Are Predictors of Insulin Resistance in Women with Gestational Diabetes Mellitus. Endocr. Metab. Immune Disord. Drug Targets 19, 358–366 (2019).

31. Kopcow, H. D. et al. Human decidual NK cells form immature activating synapses and are not cytotoxic. Proc. Natl. Acad. Sci. U. S. A. 102, 15563–15568 (2005).

32. Williams, P. J., Searle, R. F., Robson, S. C., Innes, B. A. & Bulmer, J. N. Decidual leucocyte populations in early to late gestation normal human pregnancy. J. Reprod. Immunol. 82, 24–31 (2009).

33. Tilburgs, T. et al. Fetal-maternal HLA-C mismatch is associated with decidual T cell activation and induction of functional T regulatory cells. J. Reprod. Immunol. 82, 148–157 (2009).

34. Powell, R. M. et al. Decidual T Cells Exhibit a Highly Differentiated Phenotype and Demonstrate Potential Fetal Specificity and a Strong Transcriptional Response to IFN. J. Immunol. Baltim. Md 1950 199, 3406–3417 (2017).

35. Yu, J. et al. Assessment of the number and function of macrophages in the placenta of gestational diabetes mellitus patients. J. Huazhong Univ. Sci. Technol. Med. Sci. Hua Zhong Ke Ji Xue Xue Bao Yi Xue Ying Wen Ban Huazhong Keji Daxue Xuebao Yixue Yingdewen Ban 33, 725– 729 (2013).

36. Melgert, B. N. et al. Pregnancy and preeclampsia affect monocyte subsets in humans and rats. PloS One 7, e45229 (2012).

37. Leijte, G. P. et al. Monocytic HLA-DR expression kinetics in septic shock patients with different pathogens, sites of infection and adverse outcomes. Crit. Care Lond. Engl. 24, 110 (2020).

38. Salvany-Celades, M. et al. Three Types of Functional Regulatory T Cells Control T Cell Responses at the Human Maternal-Fetal Interface. Cell Rep. 27, 2537–2547.e5 (2019).

39. Wienke, J. et al. Human Tregs at the materno-fetal interface show site-specific adaptation reminiscent of tumor Tregs. JCI Insight 5, e137926 (2020).

40. Jiang, X. et al. Role of the tumor microenvironment in PD-L1/PD-1-mediated tumor immune escape. Mol. Cancer 18, 10 (2019).

41. Francisco, L. M. et al. PD-L1 regulates the development, maintenance, and function of induced regulatory T cells. J. Exp. Med. 206, 3015–3029 (2009).

42. Lin, C.-L., Huang, H.-M., Hsieh, C.-L., Fan, C.-K. & Lee, Y.-L. Jagged1-expressing adenovirus-infected dendritic cells induce expansion of Foxp3+ regulatory T cells and alleviate T helper type 2-mediated allergic asthma in mice. Immunology 156, 199–212 (2019).

43. Zhang, A. L., Wang, F., Chang, L.-S., McDonnell, M. E. & Min, L. Coexistence of Immune Checkpoint Inhibitor-Induced Autoimmune Diabetes and Pancreatitis. Front. Endocrinol. 12, 620522 (2021).

44. Martinov, T., Spanier, J. A., Pauken, K. E. & Fife, B. T. PD-1 pathway-mediated regulation of islet-specific CD4+ T cell subsets in autoimmune diabetes. Immunoendocrinology Houst. Tex 3, e1164 (2016).

45. Zhang, Y. et al. The role of the PD-1/PD-L1 axis in macrophage differentiation and function during pregnancy. Hum. Reprod. 34, 25–36 (2019).

46. Okuyama, M., Mezawa, H., Kawai, T. & Urashima, M. Elevated Soluble PD-L1 in Pregnant Women’s Serum Suppresses the Immune Reaction. Front. Immunol. 10, 86 (2019).

47. Habicht, A. et al. A Link between PDL1 and T Regulatory Cells in Fetomaternal Tolerance. J. Immunol. 179, 5211–5219 (2007).

48. Tian, M. et al. The PD-1/PD-L1 inhibitory pathway is altered in pre-eclampsia and regulates T cell responses in pre-eclamptic rats. Sci. Rep. 6, 27683 (2016).

49. Paolino, M. et al. RANK links thymic regulatory T cells to fetal loss and gestational diabetes in pregnancy. Nature 589, 442–447 (2021).

50. Zhao, Y. et al. Immune checkpoint molecules on T cell subsets of pregnancies with preeclampsia and gestational diabetes mellitus. J. Reprod. Immunol. 142, 103208 (2020).

51. WHO recommendation on the diagnosis of gestational diabetes in pregnancy (https://srhr.org/rhl/article/who-recommendation-on-the-diagnosis-of-gestational-diabetes-in-pregnancy) Accessed on 12.10.2022 13:00.

52. Kargl, J. et al. Neutrophils dominate the immune cell composition in non-small cell lung cancer. Nat. Commun. 8, 14381 (2017).

53. Xu, Y., Plazyo, O., Romero, R., Hassan, S. S. & Gomez-Lopez, N. Isolation of Leukocytes from the Human Maternal-fetal Interface. J. Vis. Exp. JoVE e52863 (2015) doi:10.3791/52863.

54. Wickham H, Averick M, Bryan J, Chang W, McGowan LD, François R, Wickham et al., (2019). Welcome to the Tidyverse. Journal of Open Source Software, 4(43), 1686.

55. Hao, Y. et al. Integrated analysis of multimodal single-cell data. Cell 184, 3573–3587.e29 (2021).

56. Davis, S. & Meltzer, P. S. GEOquery: a bridge between the Gene Expression Omnibus (GEO) and BioConductor. Bioinforma. Oxf. Engl. 23, 1846–1847 (2007).

