## Supplementary Material for "Gestational diabetes mellitus dysregulates the PD-1/PD-L1 axis at the feto-maternal interface"

### Supplementary Fig 1

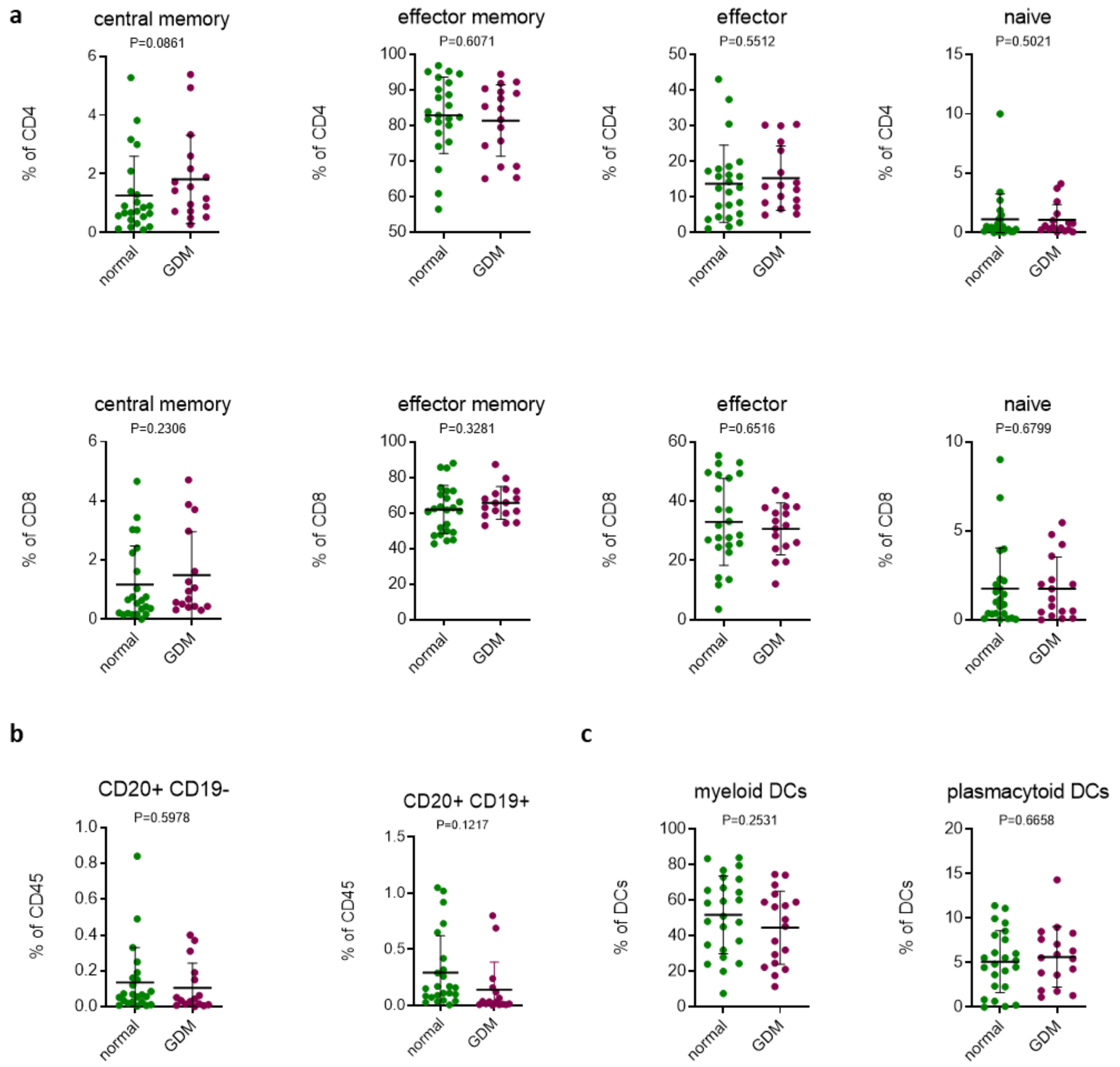


**Supplementary Figure 1. Changes in immune subpopulations between normal and GDM DB samples.**

(**a**) Abundance and changes in CD4+ and CD8+ immune populations between normal (N = 24, left) and GDM (N = 18, right) DB samples. (**b**) Abundance and changes in B cell subsets based on the expression of CD20 and CD19 between normal (N = 24, left) and GDM (N = 18, right) DB samples. (**c**) Abundance and changes DC subpopulations between normal (N = 24, left) and GDM (N = 18, right) DB samples. Mann–Whitney U tests were performed to compare two groups. Data are represented as mean ± SD, *P < 0.05.

Supplementary Fig 2

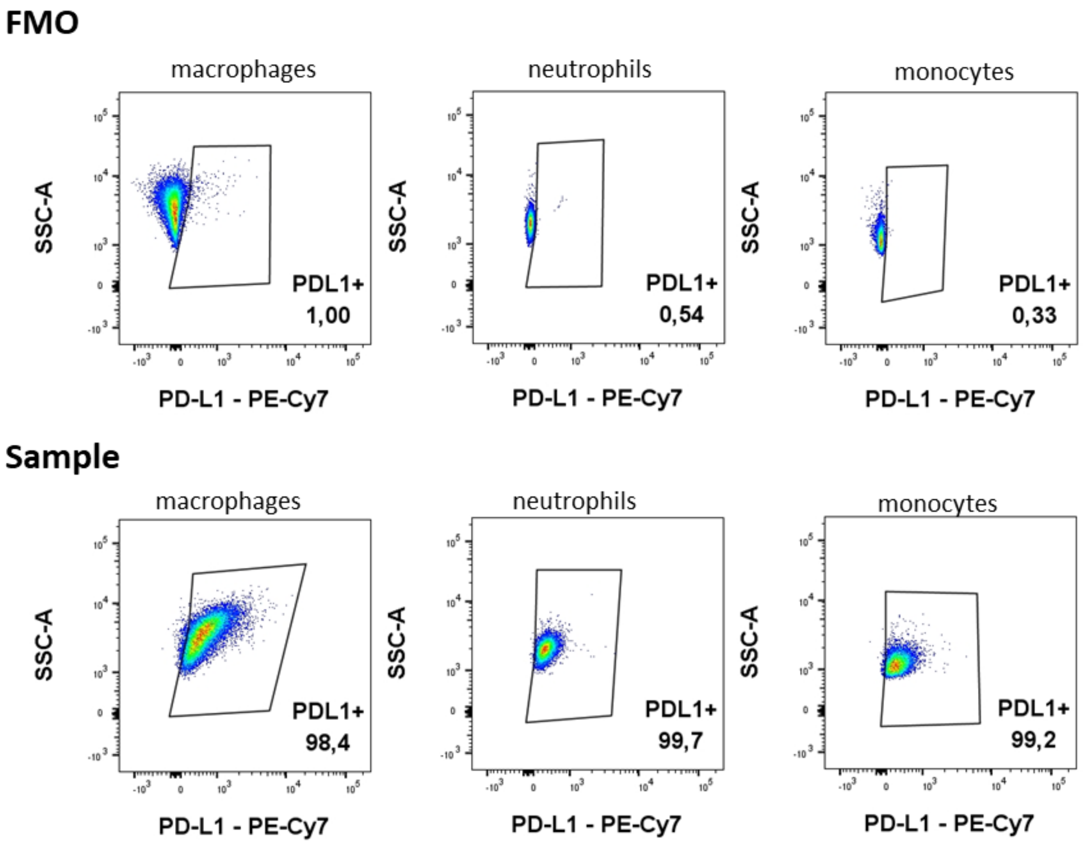


**Supplementary Figure 2. Representative flow cytometry dotplots of PD-L1 gating based on FMO**

Supplementary Fig 3

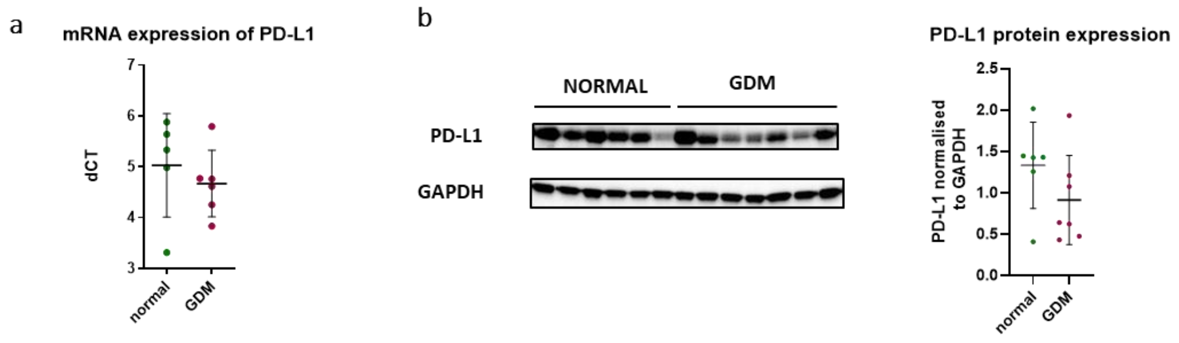


**Supplementary Figure 3. Total protein expression of PD-L1**

(**a**) mRNA expression of *PD-L1* in normal (N = 5, left) and GDM (N = 6, right) tissue homogenates. (**b**) Total PD-L1 protein expression in normal (N = 6, left) and GDM (N = 7, right) tissue homogenates. Mann–Whitney U tests were performed to compare two groups. Data are represented as mean ± SD, *P < 0.05.

### Supplementary Fig 4

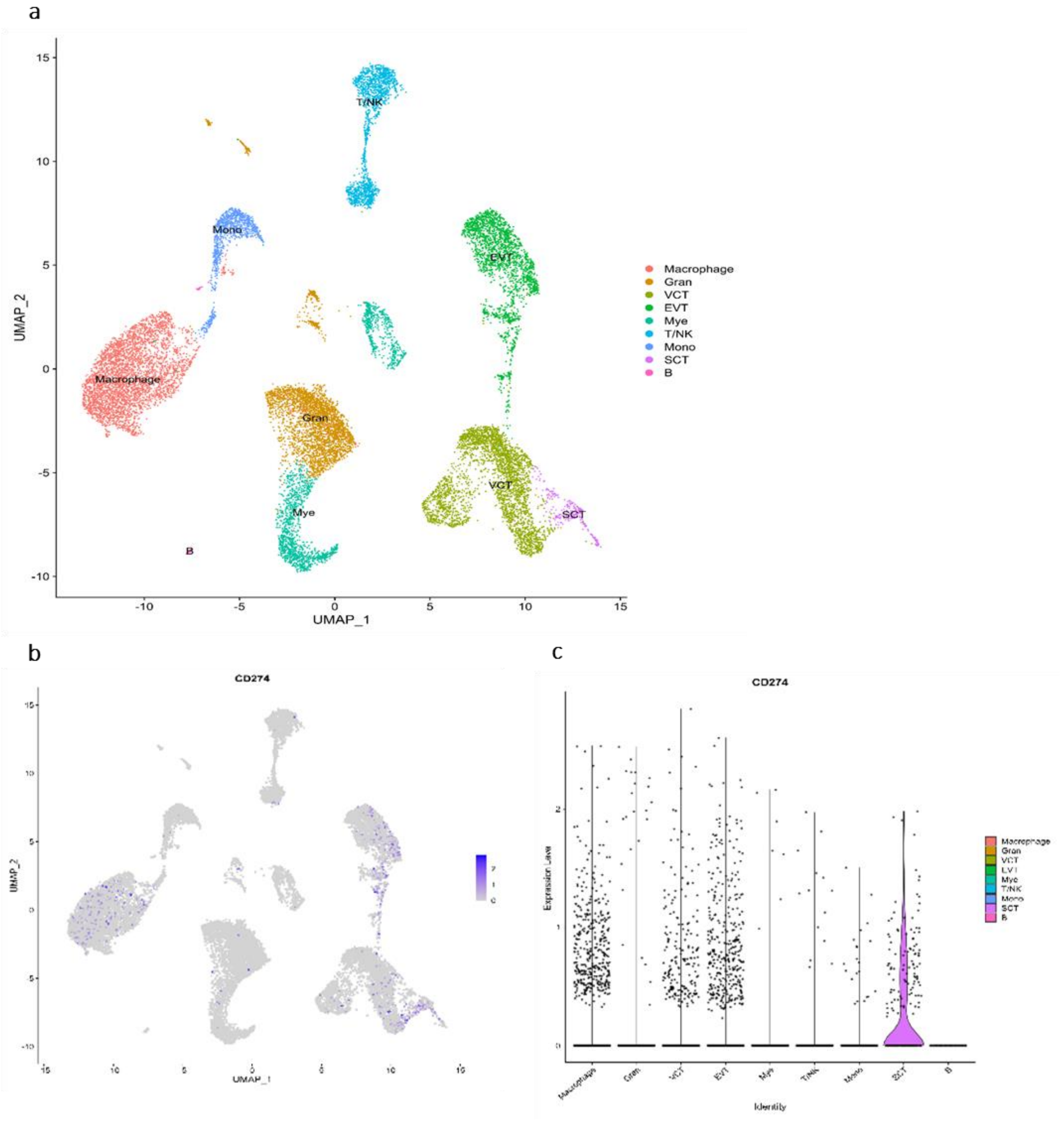


**Supplementary Figure 4. scRNAseq analysis of *PD-L1* expression in healthy and GDM DB samples.** (**a**) Uniform Manifold Approximation and Projection (UMAP) of cells infiltrating the DB. (**b**) *PD-L1* expression on the cell populations. (**c**) Abundance of *PD-L1* expression on the cell populations.

### Supplementary Fig 5

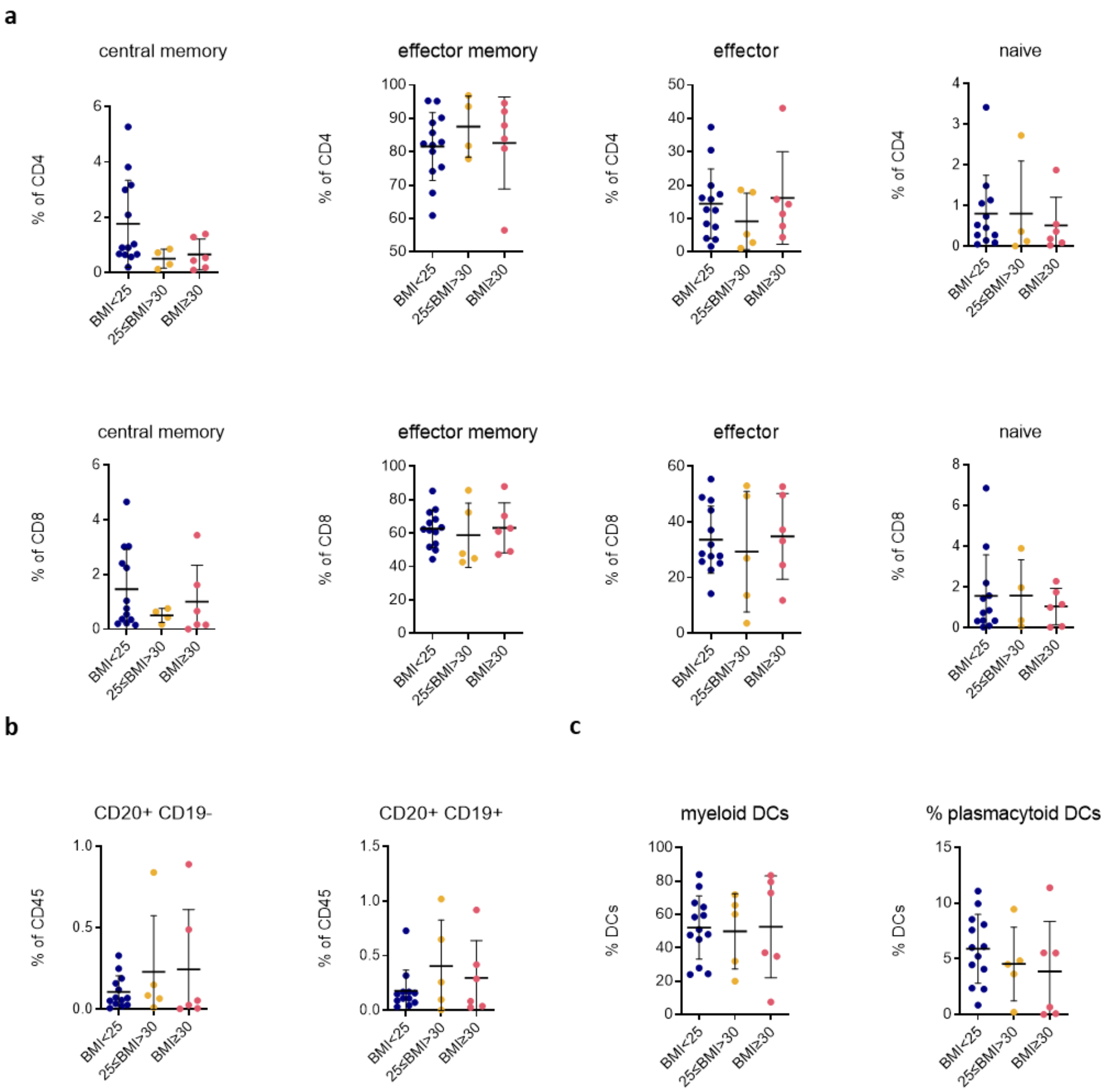


**Supplementary Figure 5. Changes in immune subpopulations between healthy donors based on BMI.**

(**a**) Abundance and changes in CD4+ and CD8+ immune populations between healthy DB samples from normal BMI, overweight, and obese mothers. (**b**) Abundance and changes in B cell subsets based on the expression of CD20 and CD19 between healthy DB samples from normal BMI, overweight, and obese mothers. (**c**) Abundance and changes in DC subpopulations between healthy DB samples from normal BMI, overweight, and obese mothers. One-way ANOVA was performed to compare three groups. Data are represented as mean ± SD, $^{*}P < 0.05$.

**Supplementary Table 1. Antibodies used for immunophenotyping**

### Supplimentary Table 1

| Marker | Fluorophore | Clone | Company |
|---|---|---|---|
| CD45 | AF700 | HI30 | BioLegend |
| CD56 | PE | HCD65 | BioLegend |
| CD16 | BV510 | 3G8 | BD |
| CD19 | BUV496 | SJ2511 | BD |
| CD20 | BUV805 | 2H7 | BD |
| CD3 | APC | UCHT1 | BioLegend |
| CD4 | BUV395 | SK3 | BD |
| CD8 | BV605 | RPA-T8 | BioLegend |
| CCR7 | PE/Dazzle594 | G043H7 | BioLegend |
| CD45RA | BV785 | HI100 | BioLegend |
| CD38 | PE-Cy7 | HB-7 | BioLegend |
| CD69 | FITC | FN50 | BioLegend |
| PD1 | BUV737 | EH12.1 | BD |
| FoxP3 | PerCpCy5.5 | 236A/E7 | BD |
| CD66b | PE | G10F5 | BioLegend |
| Siglec8 | PerCpCy5.5 | 7C9 | BioLegend |
| CD68 | FITC | Y1/82A | BioLegend |
| CD80 | BV711 | 2D10 | BioLegend |
| CD206 | APC | 15-2 | BioLegend |
| CD14 | BUV737 | M5E2 | BD |
| HLA-DR | BV510 | L234 | BioLegend |
| CD16 | BUV395 | 3G8 | BD |
| CD11c | BV605 | 3.9 | BioLegend |
| CD123 | BV421 | GH6 | BioLegend |
| PD-L1 | PE-Cy7 | B7-H1 | BioLegend |
